# DNA-based long-lived reaction-diffusion patterning in a host hydrogel

**DOI:** 10.1101/767608

**Authors:** Georg Urtel, André Estevez-Torres, Jean-Christophe Galas

## Abstract

The development of living organisms is a source of inspiration for the creation of synthetic life-like materials. Embryo development is divided into three stages that are inextricably linked: patterning, differentiation and growth. During patterning, sustained out-of-equilibrium molecular programs interpret underlying molecular cues to create well-defined concentration profiles. Implementing this patterning stage in an autonomous synthetic material is a challenge that at least requires a programmable and long-lasting out-of-equilibrium chemistry compatible with a host material. Here we show that DNA/enzyme reactions can create reaction-diffusion patterns that are extraordinary long-lasting both in solution and inside an autonomous hydrogel. The life-time and stability of these patterns - here traveling fronts and two-band patterns - are significantly increased by blocking parasitic side reactions and by dramatically reducing the diffusion coefficient of specific DNA sequences. Immersed in oil, hydrogels pattern autonomously with limited evaporation, but can also exchange chemical information from other gels when brought in contact. Our primitive metabolic material thus recapitulates two important properties of living matter: a certain degree of autonomy that makes each piece of material an ‘individual’ with its own metabolism and, at the same time, the capacity to interact with other ‘individuals’.

## Introduction

Living organisms have been a continuous source of inspiration for the development of new materials with diverse properties.^1^ From wood fibers to butterfly wings, the structure of living materials has been successfully imitated in fields as diverse as construction^2^ or optics.^3^ In addition to its precious structural qualities, living matter is also interesting due to its highly dynamic nature. The complex and sustained network of biochemical reactions - i.e. the metabolism - that links the thousands of elements that make up a biological cell brings out surprising properties. Recently, the idea of transposing the out-of-equilibrium state of living matter into synthetic materials has emerged.^4–9^ This unique property of living organisms would give synthetic metabolic materials the ability to self-construct, interact with the environment or reconfigure. ^10^

The implementation of such metabolic materials requires a tangible and autonomous piece of matter in which programmable and out-of-equilibrium chemical reaction networks take place. Historically, a hydrogel embedded with the Belousov-Zhabotinsky (BZ) oscillating reaction that shows chemomecanical transduction in response to chemical dynamics has been the very first example of such an autonomous metabolic material, and still remains unsurpassed.^11–13^ However, the low programmability and the harsh reaction conditions of the BZ reaction severely limit the developments of metabolic materials.

Providing a higher degree of programmability and biocompatibility, DNA-based molecular programs^14–16^ offer an attractive alternative to build next generation metabolic materials. In this regard, the pattern-forming mechanisms occurring during early embryo development,^17,18^ are a source of inspiration in the design of metabolic materials that are capable of spatial differentiation. ^6^ Reaction-diffusion (RD) patterning is a major mechanism of chemical differentiation,^19^ and an increasing number of RD patterns have been recently implemented with DNA-based molecular programs.^6,20–26^ However, these programmable patterns have not yet been obtained inside an autonomous tangible material. One reason for this is that complex RD patterns need the underlying chemical network to be maintained out-of-equilibrium, which in most cases implies using an open reactor, i.e. a reservoir that continuously replenishes the reaction with fresh chemical fuel.

To circumvent this problem, we take advantage of the PEN (Polymerase Exonuclease Nicking enzyme) DNA toolbox, a powerful molecular programming approach relying on short DNA oligonucleotides and enzymes, that has enabled the experimental implementation of complex temporal^15,27–29^ and spatio-temporal concentration patterns^6,21,30,31^ in closed reactors.

We first significantly extend the life-time of the RD patterns - from 5 to 60 h in the current experimental conditions - by suppressing the autocatalytic formation of parasitic strands. Later, we increase the stability of two-band patterns arising from the interpretation of an underlying morphogen gradient. To do so we achieve a 35-fold reduction of the diffusion coefficient of a reactive DNA strand by co-polymerizing it with linear polyacrylamide. ^32^

Finally, we embed this active solution inside a manipulable passive agarose hydrogel, thus creating an autonomous metabolic material capable of self-patterning. In particular, we obtained traveling fronts and immobile band patterns inside hydrogels immersed in oil. Notably, the hydrogels can be moved, deformed or put in contact by an external action without perturbing the patterning process.

## Results and discussion

### A typical reaction-diffusion system for static pattern generation

Among the non-trivial reaction-diffusion patterns that were implemented using the PEN DNA toolbox, the static two-band pattern generated from the reading of an underlying gradient is particularly interesting in the scope of a future metabolic material. Indeed, this out-of-equilibrium patterning is the simplest one capable of spatial differentiation, in a one-dimensional reactor.^6^ The position of the border in between the two regions is continuously computed from the underlying molecular gradient.

To this end, we use a bistable network that is able to bifurcate from the production of a given DNA strand **A** to the destruction of the same strand depending on the concentration of another DNA species, the repressor **R**.^33^ As depicted in Figure 1a and b and more detailed in Figure S1, the reaction network is based on three reactions: (i) the autocatalytic production of **A**, a 12-mer single stranded DNA (ssDNA), supported by template **T**, a 22-mer ssDNA, (ii) the repression of **A** by **R**, a 16-mer ssDNA that produces the waste **W**, a 16-mer ssDNA, and (iii) the degradation of **A** and **W**. DNA sequences are detailed in Table S1. Three enzymes are involved in the reaction: a polymerase (pol), a nicking enzyme (nick) and an exonuclease (exo). **T** and **R** are protected against degradation by the exonuclease.

**Figure 1:**
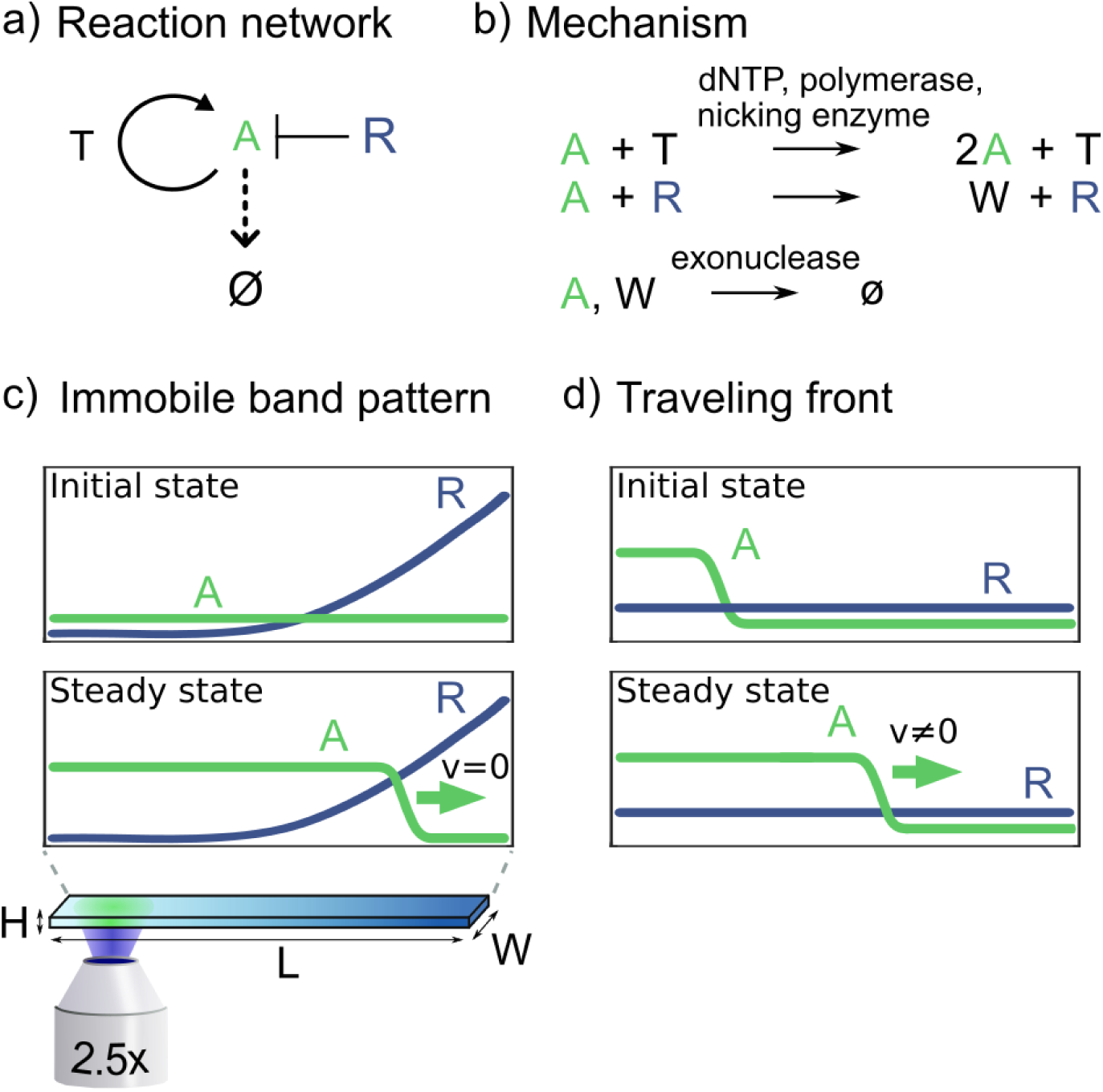
A bistable DNA-based reaction network generates an immobile band pattern or a traveling front. a) The bistable reaction network consists of an autocatalytic reaction of **A** supported by template **T**, a repression of **A** by **R** and a degradation of **A**. b) Details of the reactions shown in panel a. c) Scheme of a two-band pattern-formation experiment: a gradient of **R** is initially created inside a capillary. The production of **A** will then initiate where the concentration of **R** is low, thus generating a front of **A**. The front will propagate towards the higher value of **R** until it reaches the bifurcation point of the bistable system and stops, forming two chemically distinct zones (high concentration of **A**, left, low concentration of **A**, right). Experimentally, a one-dimensional system was used with the reactor length *L* much greater than the reactor width *W* and height *H*. d) Scheme of a traveling front experiment: when the autocatalytic production of **A** dominates over the repression of **A** by **R**, an initial excess of **A** on one side of the capillary gives rise to a traveling front that never stops.

The system is designed in a way that the repression of **A** by **R** occurs faster than the production of **A** by **T**. The concentration of **R** is a bifurcation parameter of the bistable network.^15^ Below a given concentration of **R**, the autocatalytic production of **A** is active and the concentration of **A** in the solution increases. Above this threshold, the autocatalytic production of **A** cannot be triggered as **A** strands are degraded into **W**, and the concentration of **A** tends to zero.

As depicted in Figure 1c, in the presence of an underlying gradient of **R** in a 1-dimensional reactor, a static pattern will form. Initially, the production of **A** will begin where the concentration of **R** is the lowest. A reaction-diffusion front will then propagate towards the highest **R** concentration, and it will continuously slow down until reaching the bifurcation point of the network. At this position, the front will stop, generating an out-of-equilibrium static two-band pattern.

This reaction network can also be used to generate a wavefront that travels at constant velocity as depicted in Figure 1d. In this case, the concentration of **R** has to be homogeneously low. It will prevent early phase background amplification - a well known problem often simply called ‘selfstart’.^33,34^ And **A** has to be introduced on one side of the channel to trigger the reaction-diffusion front.

### Concentration gradients were created both in solution and inside hydrogels, and observed with limited evaporation

A two-band pattern spatio-temporal experiment requires a gradient of repressor **R** and a homogeneous concentration of all other species. We developed a simple protocol to create a gradient of **R** by Taylor dispersion through back and forth pipetting inside a capillary. The capillary is first filled with a solution containing the DNA strands and the enzymes but not **R**. Then, the open end of the capillary is immersed in a solution containing **R**, and back and forth pipetting is performed, which results in the mixing of the two solutions by Taylor dispersion and the creation of a smooth gradient of **R** along the longitudinal axis of the capillary as shown in Figure 2a. While manual, the method is highly reproducible (Figure 2b). It can also be automated using a programmable syringe pump for convenience, but reproducibility was not significantly improved (Figure S4 and Movie S1).

**Figure 2:**
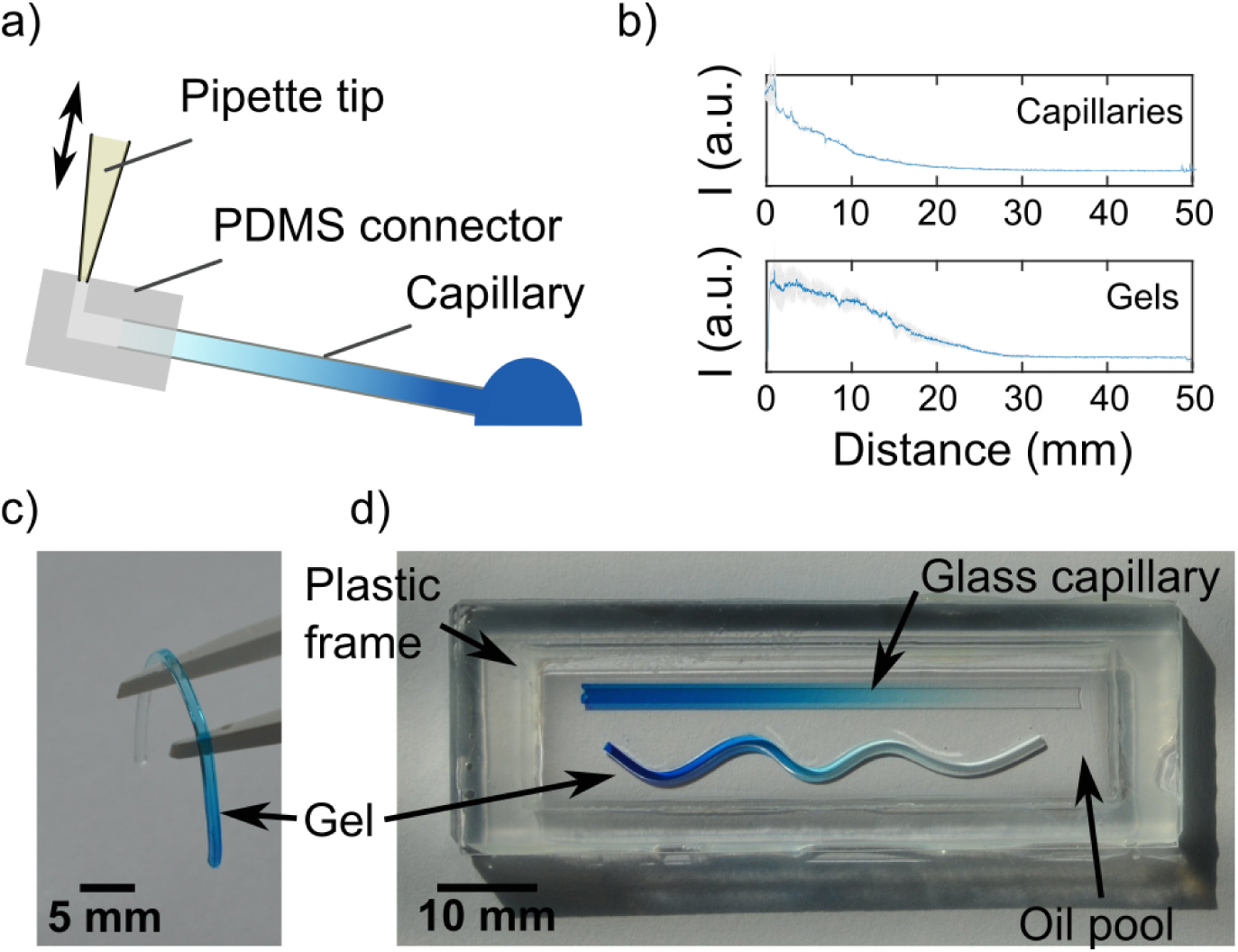
Centimeter-sized gradients were reproducibly generated both in an aqueous solution and in a hydrogel. a) Scheme of the gradient-generation protocol. A low concentration solution (light blue) inside a glass capillary is brought in contact with a droplet of high concentrated solution (dark blue). Subsequently, a connector is used to perform back and forth pipetting and create a gradient along the channel length through Taylor dispersion. b) Fluorescence profiles of 22-mer DNA gradients obtained inside capillaries (top panel) or gels (bottom panel). Blue lines represent the mean of 3 profiles, standard deviation appears in gray. c) A piece of agarose gel with an embedded gradient. d) Two gradients (inside a 0.2×2×50 mm glass capillary and in a 1×1×50 mm gel) immersed into a pool of mineral oil to keep the temperature constant and avoid evaporation during long experiments.

The same protocol was used to form a tube of agarose gel with an embedded gradient. In this case, the capillary and the solutions were heated at 60°C to keep the gel liquid during the gradient preparation. At this temperature, the enzyme activity is not irreversibly affected (Figure S5). After cooling, the gel was extracted from the capillary by pushing it outwards. A 1 × 1 × 50 mm gel tube that can be easily manipulated and which contains a methylene blue dye gradient is shown Figure 2c. The reusable polydimethylsiloxane connector avoids the use of a microfluidic gel molding device and makes the method very simple.^35^

Pattern formation in our experiments occurs typically at 44°C, which induces important evaporation problems and preclude long-term experiments. In previous works, we were sealing the capillary ends with glue or grease. ^6,36^ However, the fluorescence of the glue or the grease prevented the ends of the capillaries from being imaged. In addition, this method cannot be applied to molded agarose gel. Here, we show that immersing the samples, capillaries or gels, into a pool of oil, as shown in Figure 2d, considerably limits evaporation. We measured that less than 1% of volume evaporates inside a capillary after 3 days at 44°C. Evaporation from the gel occurs through a large surface and is thus higher, we measured a 14% volume contraction over 24 hours (Figures S7 and S8). Otherwise, the oil favors a homogeneous temperature, and the quality of fluorescence imaging is maintained over the entire length of the samples.

### Long-lived traveling fronts and band patterns were obtained by suppressing the formation of parasitic autocatalytic strands

The PEN DNA toolbox relies on the activity of enzymes that produce or degrade the DNA strands involved in the designed reaction network. While those enzymes can keep the system out-of-equilibrium in a closed reactor by consuming dNTPs, long term experiments are disrupted by a well known phenomena: the appearance of long strands of unwanted DNA called ‘parasite’.^34,37^ These DNA strands of various lengths correspond to sequences synthesized de novo by the DNA polymerase. They contain the nicking enzyme recognition site and are thus exponentially amplified. They quickly saturate the active solution, consuming the chemical fuel, sequestering the enzymatic machinery and disrupting the reaction-diffusion patterns. In a recent work, we have proposed a method to prevent the formation of parasite strands, by suppressing untemplated replication in the exponential amplification reaction (EXPAR) present in PEN DNA toolbox reactions.^36^ The method relies on specific nicking enzymes, such as Nb.BssSI, that bear only three types of nucleobases on each of the strands of the recognition site (Figure S6). In that case, fully functional replication templates can be designed using A, C, and G nucleobases only, that will produce DNA strands containing, respectively, T, G, and C nucleobases only (Figure S6). In the absence of dATP in the buffer, any emerging parasitic strand will not contain the A nucleobase. It will, therefore, not carry the nicking enzyme recognition site and will not be replicated exponentially, unlike the designed strand **A**.

We successfully applied this method on the network presented above for reaction-diffusion patterning. Strands **T** and **R** only contain the A, C, and G nucleobases and the reaction buffer only contains dTTP, dCTP and dGTP. The left kymographs of Figure 3a and b show a reaction-diffusion front experiment and a two-band pattern experiment in the presence of all four dNTPs. These experiments are performed in a capillary filled with an aqueous solution that contains the intercalating dye EvaGreen to monitor the dsDNA concentration over space and time (Figure S9). Experiments last for 17 hours, but after 2 hours several highly fluorescent bumps indicate the emergence of a growing parasite that disrupt the expected patterns. In contrast, in the absence of dATP, no parasitic strand production is observed and the traveling front and the two-band pattern are sustained throughout the experiment (right kymographs in Figure 3a and b and Figure S10 for concentration profiles). In the following, we will take advantage of this method to perform experiments that last for more than 60 hours.

**Figure 3:**
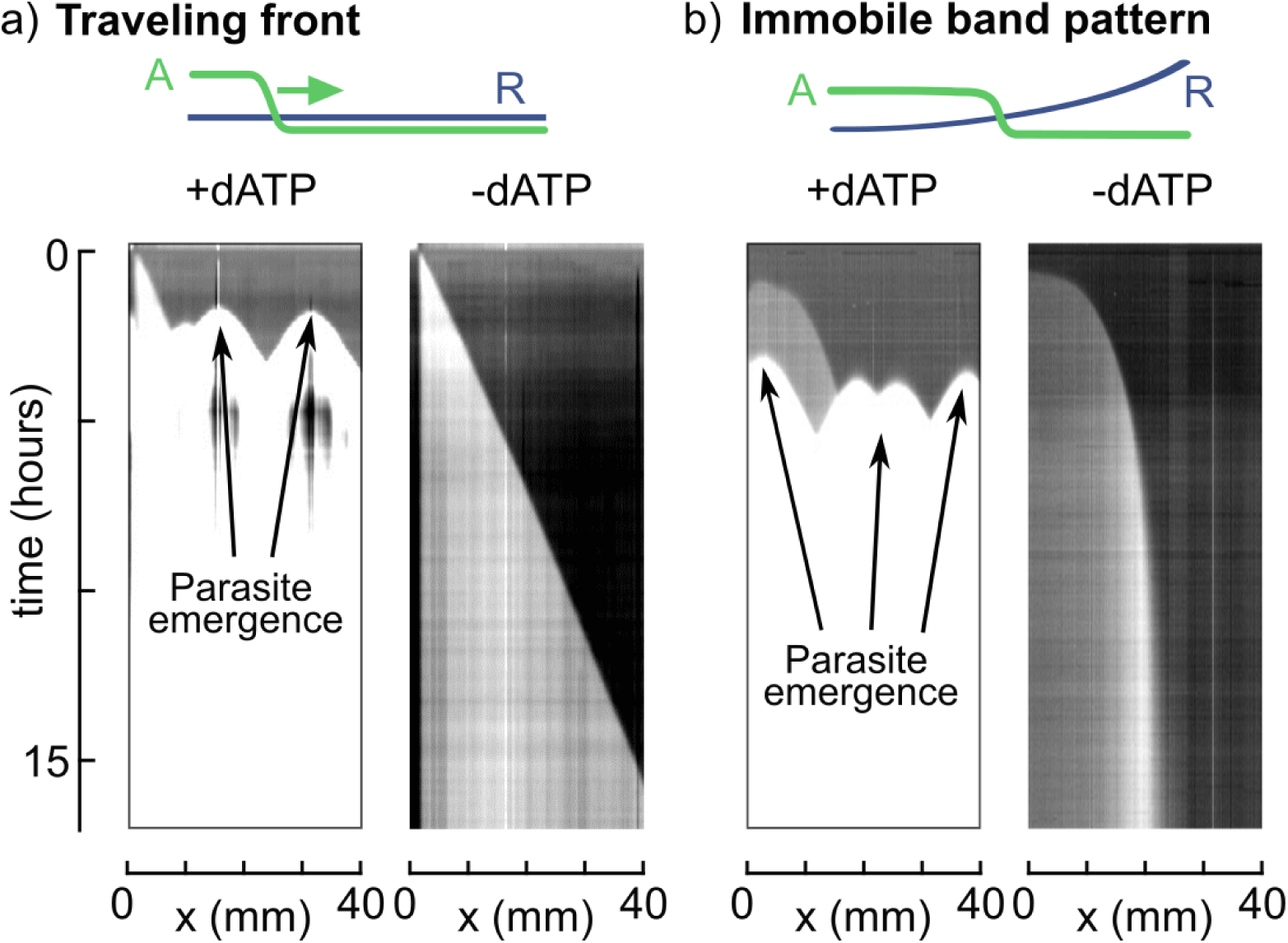
Parasite suppression allows long lived reaction-diffusion patterns. a) Kymographs (time vs position, fluorescence shift due to high concentration of **A**) of traveling fronts. Fronts are initiated on the left side of the capillaries and propagate with uniform velocity. In the presence of dATP (left) parasites emerge after 2 hours at different positions in the capillary. In the absence of dATP (right) parasites never emerge and the front travels through the whole capillary in 15 hours. b) Kymographs of two-band patterns. In the presence of dATP (left) parasites emerge after 5 hours at different positions in the capillary. In the absence of dATP (right) parasites never emerge, the front travels from left to righ progressively decelerating until it stops, forming a two-band pattern with high concentration of **A** on the left and absence of **A** on the right. For all experiments, conditions favoring parasite emergence were used. Conditions, traveling front: 32 U/mL pol, 200 U/mL nick, 2% exo, 50 nM **T**, 50 nM **R**, and 0.4 mM dNTPs with or without dATP. Front is triggered using 0.5 *µ*L of 1 *µ*M **A**. Conditions, immobile band pattern: 32 U/mL pol, 200 U/mL nick, 2% exo, 50 nM **T**, 0-1000 nM **R**, and 0.4 mM dNTPs with or without dATP.

### Decoupling the dynamics of reactive species enables the preservation of underlying molecular information

Taking advantage of the absence of parasite, the left kymograph of Figure 4a shows an experiment that lasts for more than 60 hours. However, as it can be seen, the boundary between the two regions of high and low DNA concentration is not stable over time. The front velocity over time (Figure 4a, bottom panel, blue plot) confirms this observation: it decreases to reach a constant velocity *v* =4.5 *µ*m/min over 20 hours before increasing after 40 hours until reaching 15 *µ*m/min.

**Figure 4:**
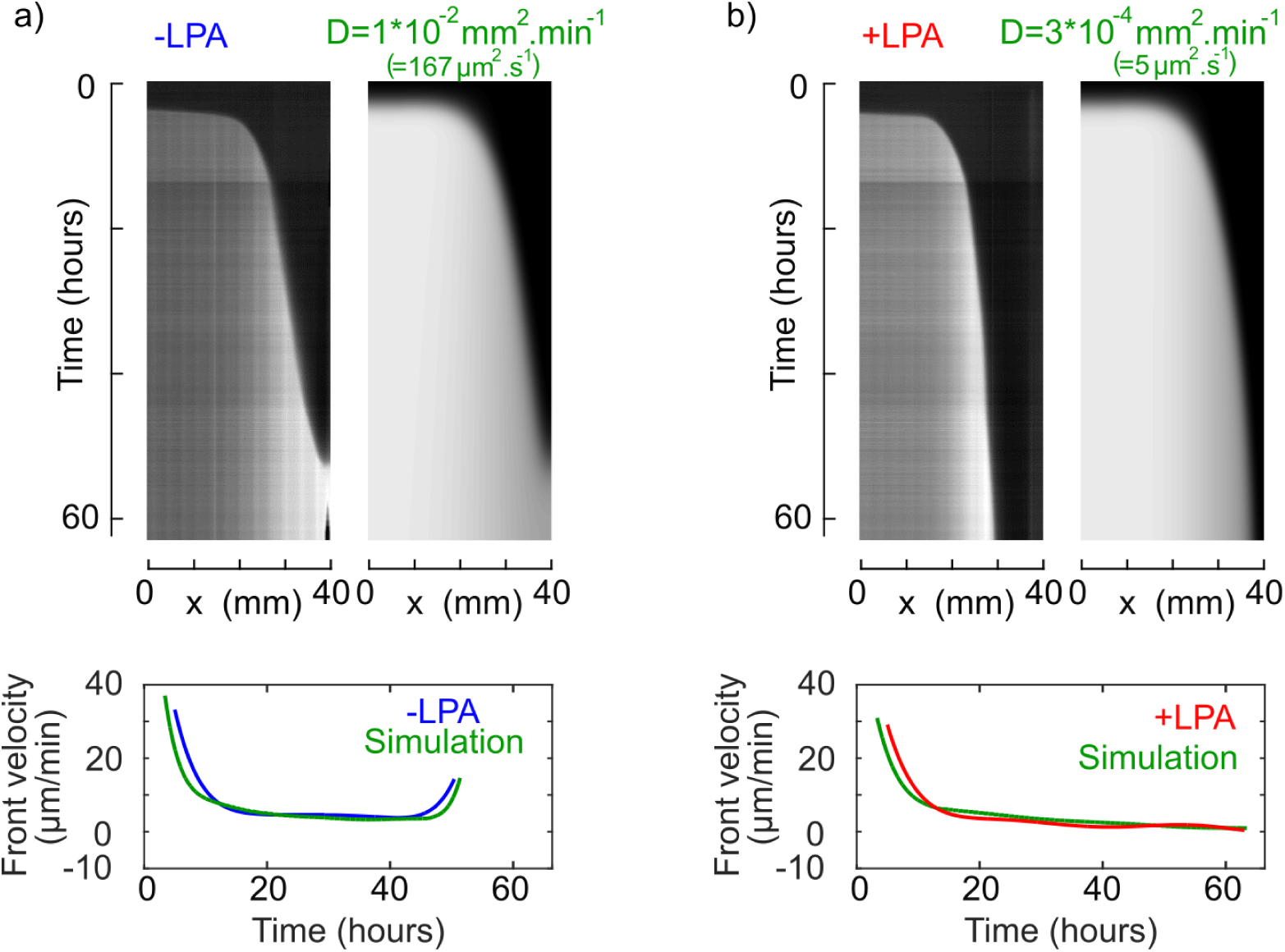
Long-term stability of the final pattern is significantly increased by a slowly diffusing underlying cue. a) Top: experimental and simulated kymographs (time vs position, fluorescence shift due to **A** strand) of a two-band pattern experiment in the presence of a gradient of repressor **R** that freely diffuses. The pattern forms and disappears as the out-of-equilibrium molecular program interprets a continuously changing gradient cue. Bottom: front velocities over time. b) Top: experimental and simulated kymographs of a two-band pattern experiment in the presence of a gradient of repressor attached to linear polyacrylamide (**LPA-R**, diffusing 35 times less). The pattern is stable for more than 60 hours. Bottom: front velocities over time. Experimental conditions: 8 U/mL pol, 200 U/mL nick, 2% exo, 50 nM **T**, 0-750 nM **R**, and 0.4 mM dNTPs. Simulation parameters are detailed in the Supporting Information section 10.

To explain the long-term destabilization of the two-band pattern, our hypothesis is that the diffusion of the underlying gradient modifies the lateral position for which the bifurcation concentration of the bistable system is reached. This concentration threshold shifts towards the right side of the capillary, making the high fluorescence region drifting to the right, which is clearly visible after 45-50 hours.

Indeed, considering the diffusion of **R** over *L* =3 mm with a diffusion coefficient *D* =170 *µ*m^2^/s, **R** will diffuse with a characteristic time *τ* _*D*_ = *L*^2^/2*D≈* 14 h. This means that the gradient of **R** will significantly flatten during long lasting experiment, potentially compromising the stability of the band pattern (Figure S11).

Numerical simulations of a simple model (Supporting Information section 10) accounting for the autocatalytic production of **A** over template **T**, the repression of **A** by **R**, the degradation reactions and the diffusion of the underlying **R** gradient qualitatively catches this phenomenon (Figure 4a).

To stabilize the two-band pattern at longer times, we decided to immobilize, or drastically reduce the diffusion of **R** in order to impede the underlying gradient to flatten. Surface attachment is a possible strategy. Yet it is difficult to obtain a pattern showing a clean gradient over several centimeters. ^38^ Moreover, the distribution of the DNA in the solution is not homogeneous, which may disrupt the performance of the reaction network.

We chose an alternative strategy by copolymerizing the DNA repressor into a long polymer chain. In particular, a DNA strand modified at its 5’ end by an acrylic phosphoramidite was incorporated into a linear polyacrylamide (LPA) chain during the free radical polymerization, thus forming a poly(acrylamide-co-DNA).^32,39^

The purification of the solution - in particular to remove the non-attached DNA or the DNA attached to short LPA chains - was carried out by electroelution. We used size-exclusion chromatography to estimate the weight distribution of the LPA-DNA conjugate, and found a molecular weight in the range 0.3-2 MDa (Supporting Information section 8).

Fluorescence Recovery After Photobleaching (FRAP) experiments were performed to determine the diffusion coefficient of the LPA-DNA conjugate at 44°C. We found *D* =4.8±2 *µ*m^2^/s, which is 35 times less than the free diffusing DNA (*D* =170±42 *µ*m^2^/s) (Figure S13). Inside an agarose gel matrix, the diffusion of the LPA-DNA conjugate is even smaller, *D* =2±1 *µ*m^2^/s.

Panel b in Figure 4 shows an experiment where a copolymer of **R** with LPA, **LPA-R**, was used to make the gradient of repressor. In this case, the band pattern is stable for more than 60 hours. The front velocity (Figure 4b, bottom panel), after an initial decrease due to the pattern formation remains constant and close to zero (*v* =1.5 *µ*m/min, to be compared with the traveling front velocity *v* =40 *µ*m/min shown in Figure 3). Simulations by only changing the diffusion coefficient of **R** to that of **LPA-R** are in agreement with this result (Figure 4b).

These experiments demonstrate that an individual control of diffusion for each species can be achieved. It is thus possible to implement reaction-diffusion mechanisms in the presence of an underlying molecular information that is stable over a long period of time. It recapitulates a key feature of living materials: they are not only hierarchically assembled from a structural point of view, but also from a dynamic point of view. A well-know example is the conceptual French flag model coming from developmental biology, where an immobile concentration gradient provides positional information and supports the formation of a band pattern of concentration.^40^

### Autonomous self-patterning hydrogels that are capable of exchanging chemical information

In the above, the designed active solutions were able to perform non-equilibrium spatiotemporal computations that generated non-trivial concentration patterns in a sealed reactor. Despite their interest, these computations did not occur inside a tangible piece of matter and they were not able to exchange chemical information with the outside world, two crucial properties of non-equilibrium computations in living systems. In the following, we seek to obtain metabolic materials that are simple yet combine a certain degree of autonomy with the capacity of interacting with other materials, as living systems do. In particular, we show that hydrogels embedded with DNA/enzyme active solutions are at the same time autonomous and can interact with each other.

First, we tested whether DNA/enzyme solutions were still active and capable of autonomous patterning inside a hydrogel. The hydrogel was submerged in oil to limit evaporation and provide an environment where hydrogels could be isolated and possibly also interact with each other (see below). Following the protocol described in Figure 2, we prepared an agarose rectangular block, with dimensions 1 × 1 × 50 mm, that contained a homogeneous mixture of ssDNAs **T** and **A** and of the three enzymes, and a concentration gradient of repressor strand **R** along the long axis of the hydrogel block. We obtained a piece of gel that, as expected, could be easily manipulated and deformed inside the oil pool as shown in Figure 2d. However, to facilitate image acquisition the gel was aligned as straight as possible. Figure 5 shows that a gel containing such an active solution was able to create the expected two-band reaction-diffusion pattern. If we compare it with the experiment performed in solution in similar conditions (Figure 3), the patterns look alike. In both cases dsDNA fluorescence appears at ∼ 40 min and the two-band pattern is stable for at least 10 h. However, the pattern stabilizes faster in the gel (after 5 h) that it does in solution (after 8 h), possibly because the gradient of **R** was sharper in the gel. Importantly, the activity of the enzymes is not degraded during the fabrication of the self-patterning hydrogel that takes place at 60°C (Figure S5). Finally, the parasite-suppression strategy in the absence of dATP is again fully functional.

**Figure 5:**
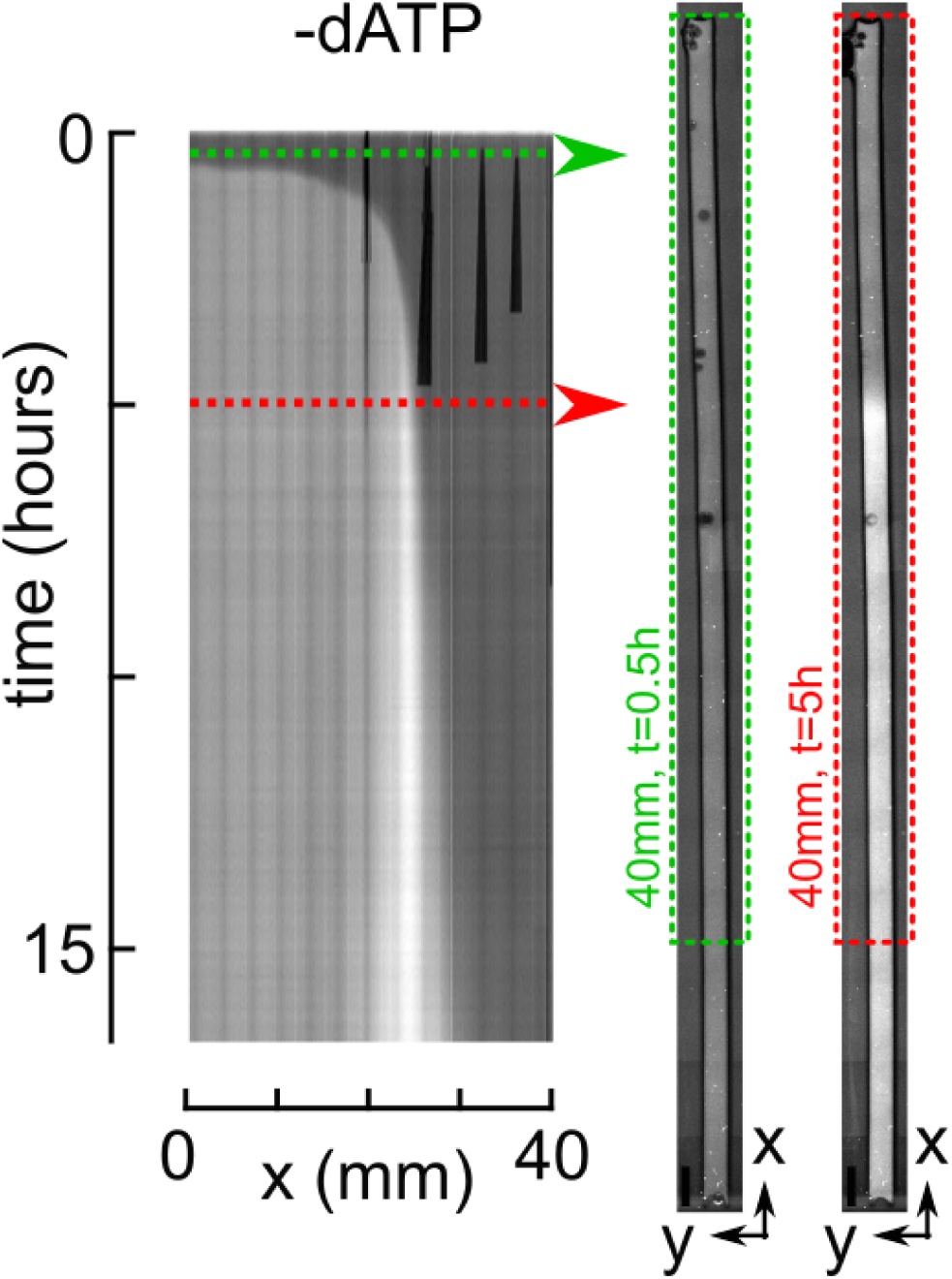
Reaction-diffusion patterning inside an autonomous hydrogel. Kymograph of the fluorescence shift due the concentration of strand **A** *vs.* time and position inside a rectangular block of gel submerged in oil. Vertical dark lines visible on top of the kymograph are air bubbles in the oil introduced when the gel was poured into the oil pool. On the right are shown fluorescent images of the full gel at *t* = 0.5 h and *t* = 5 h, imaged from the bottom. The block of gel is 1 ×1 ×50 mm^3^. Experimental conditions: 44°C, 32 U/mL pol, 200 U/mL nick, 1% exo, 50 nM **T**, 0-1000 nM **R**, and 0.4 mM dNTPs.

Second, we tested whether isolated pieces of active hydrogel could interact with each other and transfer chemical information. To do so, we fabricated three pieces of gel. One piece was a 25 mm-long rectangular block embedded with an active solution programmed to sustain a traveling wavefront but not to trigger it in the absence of an external stimulus (homogeneous concentrations of **T, R** and enzymes, but no **A**). The other two pieces were 2 mm-long agarose blocks containing the same solution supplemented with 100 nM **A**, and thus capable of triggering a traveling front of **A**. The three pieces were submerged in oil and separated from one another. At *t* = 0 min we brought into contact one small piece with the long block, and observed that a traveling front was triggered inside the long block (Figure 6). Later, at *t* = 130 min, the second small piece of gel was put in contact with the long block at a different position, triggering a new front. As expected, all fronts propagate at the same velocity and the two fronts that traveled in opposite directions annihilated at collision. ^30^ Importantly, the oil environment allowed to isolate the hydrogels when they were not in contact and to couple them when they touched each other.

**Figure 6:**
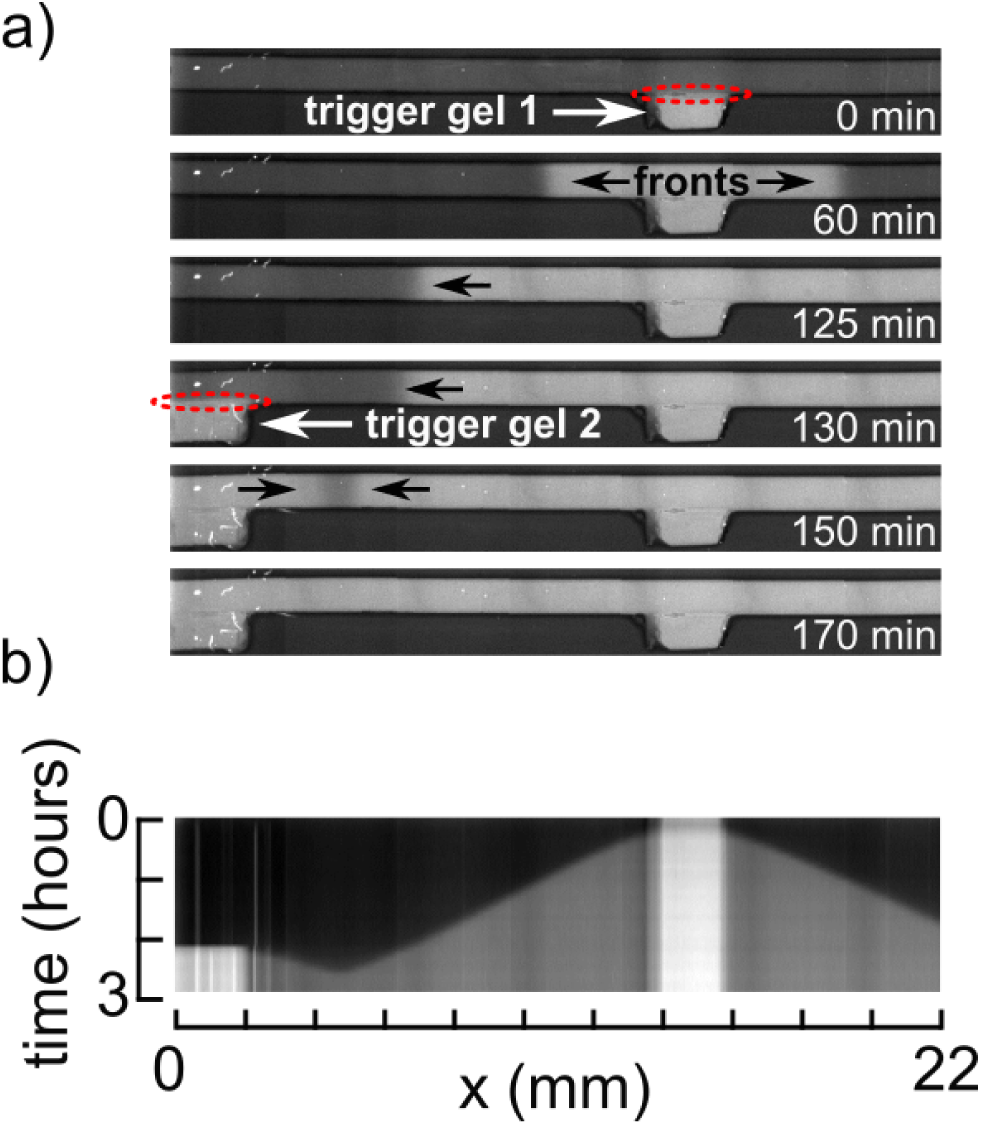
Autonomous hydrogels exchange chemical information when brought into contact. Time-lapse fluorescent images (a) and kymograph of the fluorescence shift due the concentration of strand **A** *vs.* time and position (b) showing the propagation of fronts triggered at different times (after 0 and 130 minutes) and positions through the contact (red dashed line circles) of small pieces of gel containing 100 nM of trigger **A**. Experimental conditions: 44°C, 32 U/mL pol, 200 U/mL nick, 1% exo, 50 nM T, 20 nM R, 0.4 mM dNTPs and an addition of 100 nM A for the trigger gels.

Taking together, these experiments show that an isolated metabolic material made of a hydrogel embedded with a DNA/enzyme active solution can sustain non-trivial out-of-equilibrium spatio-temporal computations over tens of hours without external input. Moreover, physical contact between different pieces of the metabolic material triggers the transfer of chemical information by diffusion. Our primitive metabolic material thus recapitulates two important properties of living matter: a certain degree of autonomy that makes each piece of material an ‘individual’ with its own metabolism and, at the same time, the capacity to interact with other ‘individuals’.

## Conclusion

Among the biomimetic materials under development, metabolic materials that combine a programmable out-of-equilibrium chemical solution coupled to a chemically responsive material, are becoming increasingly important.^10^ They seek to mimic what gives life its unique character: a sustained dynamic state, which offers self-construction and self-reconfiguration capabilities and allows constant interaction with the environment.

We took advantage of DNA/enzyme reactions to mimic key features of living systems that we would like to see incorporated into synthetic metabolic materials. To do so we solved three important limitations: (i) long-term sustainability of a well defined non-equilibrium state, (ii) dynamic heterogeneity of reaction network components in solution and (iii) autonomy and communication capability inside a tangible material.

Regarding the long term sustainability, the main two issues were the production of parasitic autocatalytic strands that saturate the samples and the diffusion of the underlying molecular gradient that disrupts the static reaction-diffusion patterns.

By using a particular nicking enzyme that contains only three types of nucleobases at the bottom strand of the restriction site (Nb.BssSI), and by limiting the dNTPs solution to dTTP, dCTP and dGTP, the formation of parasitic autocatalytic strands was thus impossible. Only voluntarily introduced templates that contain the A nucleobase necessary for nicking were able to generate exponential growth. Otherwise, to maintain underlying spatial molecular information over long time scales, we copolymerized a specific DNA strand with a long inert polymer to create a reactive species with very limited diffusion. As a result, 60 hour-long out-of-equilibrium patterns were obtained.

Finally, we embedded our active solution in a host hydrogel to get a tangible, soft and shapable autonomous metabolic material that supports non-trivial pattern formation. Interaction between gels of different compositions was demonstrated. Considering a bistable reaction network, we generated an out-of-equilibrium two-band pattern by reading an immobile underlying gradient of DNA prepared using a simple but robust experimental protocol. From a shallow gradient, the 50 mm long gel splits into two chemically distinct zones that could be used in the future to define the material fate.

The strategy demonstrated here is thus a great opportunity to explore in the near future chemomechanical transduction in a programmable metabolic material by using currently existing DNA-based chemomechanical transduction schemes.^41,42^ It provides key developments towards autonomous metabolic materials that could imitate in an ever more faithful way the extraordinary properties of living matter.

## Methods

### DNA solutions

Oligonucleotides were purchased from IDT or biomers.net (Sequences are displayed Table S1). The enzymes we used were 8-32 U/mL Bst DNA Polymerase, Large Fragment (NEB), 200 U/ml Nb.BssSI (NEB) and in-house produced *thermus thermophilus* RecJ exonuclease.^43^ 1% exonuclease corresponds to a degradation rate of 22 nM/min for a 15-mer ssDNA at 44°C. The reaction buffer contained 20 mM Tris-HCl, 10 mM (NH_4_)_2_SO_4_, 50 mM NaCl, 1 mM KCl, 2 mM MgSO4, 6 mM MgCl_2_, 1 g/L synperonic F 108 (Sigma-Aldrich), 4 mM dithiothreitol, 0.1 g/L BSA (NEB) and 1x EvaGreen Dye (Biotium). Deoxynucleotide triphosphates were purchased from NEB or Invitrogen and added in different concentrations, as specified. Kinetic characterisation experiments were performed in a CFX96 Touch Real-Time PCR Detection System (Bio-Rad) or a Qiagen Rotor-Gene qPCR machine. Epifluorescence microscopy was used for spatio-temporal experiments.

### DNA-LPA synthesis

To synthesize the DNA-linear polyacrylamide (LPA) copolymer, we purchased DNA with a 5’-acrydite modification. Monomeric acrylamide (20% w/w, Sigma-Aldrich), tetramethylethylenediamine (0.12% w/v TEMED, Sigma-Aldrich) and ammonium persulfate (0.12% w/v APS, Sigma-Aldrich) were used to polymerize linear acrylamide in the presence of 30 *µ*M acrydite-modified DNA, thus producing poly(acrylamide-co-DNA). Unreacted acrydite strands or strands with only small LPA attachements were removed by electroelution. For electroelution, we used a lab-built device, with a 5 mm wide central chamber that is sandwiched between two porous membranes with 0.22 *µ*m diameter holes. 98 *µ*L of the LPA-DNA solution was mixed with 2 *µ*L 50×TAE buffer and filled into the chamber. Each membrane separate the chamber from a tank containing 3.5 mL 1×TAE buffer. We put an electrode into each tank and ran a current of 3 mA, resulting in a voltage of approximately 50-80 V, for 10 minutes. We then recovered the sample from the central chamber. Fluorescence Recovery After Photobleaching (FRAP) and FFT image analysis using Matlab were used to extract the diffusion coefficient of the DNA-LPA.

### Experiment setup and gradient preparation

To create a gradient of DNA strand **R** in a homogeneous solution of all other species, two solutions were prepared, one with **R** and the other without. Both solutions contained the buffer, the enzymes and the rest of the DNA species. A 0.2 × 2 × 50 mm rectangular glass capillary (Vitrocom), inserted in a PDMS connector (as depicted in Figure 2), was first filled with the solution containing no **R**. Then, the end of the capillary was dipped into the reaction solution containing **R**. Micropipette up-and-down pumps of 5 to 10 *µ*L were performed to create the gradient by Taylor dispersion. After 10 to 20 back and forth pumps the gradient was formed, the connector removed and the capillary was then dropped into a pool of mineral oil (Sigma) made by a plastic frame stuck on a glass slide. The same protocol was used to mold low gelling agarose (Sigma A9539, low EEO) with embedded DNA/enzyme solutions. It has to be performed between 50°C and 70°C to get liquid agarose and avoid enzyme degradation. 1 × 1 × 50 mm capillaries were used to finally get pieces of gel that can be easily handled by hand. Once prepared, the gel was cooled down in the fridge and extracted from the capillary by pushing it outwards with a suitable piston. Again, the piece of agarose was dropped into a pool of mineral oil to avoid evaporation. The temperature of the samples, capillaries or gels immersed in the oil pool, was controlled with a Tokai Hit heating stage set at 44°C.

### Microscopy imaging

DNA concentration over space and time was measured by fluorescence. A Zeiss Axio Observer Z1 fully automated epifluorescence microscope controlled with MicroManager 1.4 was used. Contiguous fluorescence images (green channel) were recorded automatically every 1 to 20 minutes to get an entire view of the capillaries or gels after stitching. Kymographs were obtained by averaging the fluorescence images over the width of the capillary or the gel and then stacking these profiles over time.

More details can be found in the supplementary materials.

## Supporting Information Available

The supplementary data contains DNA sequences, method for gradient generation, protocol to synthesize DNA attached to LPA, characterization of DNA-LPA strand, and supplementary experiments associated to Figures 3, 4, 5 and 6.

## Acknowledgement

A. Shoushtarizadeh for preliminary experiments on gradient generation. G. McCallum for work on simulations. E. Edeleva, A. Senoussi, M.Van Der Hofstadt, Y. Rondelez and G. Ginés for insightful discussions and comments.

