## Supplementary Material for "DNA-based long-lived reaction-diffusion patterning in a host hydrogel"

### Supplementary information for: DNA-based long-lived reaction-diffusion patterning in a host hydrogel

**2 Stability of the enzymes after temperature perturbation** **12**

**3 Parasite suppression strategy** **14**

**4 Quantification of evaporation** **15**

**5 Parasite in 1D** **17**

**6 Front concentration profiles** **18**

**7 Underlying gradient diffusion with and without LPA** **19**

**8 Estimation of the molecular weight distribution of the LPA-DNA conjugate** **20**

8.1 By size exclusion chormatography . . . . . . . . . . . . . . . . . . . . 20

8.2 From the diffusion coefficient . . . . . . . . . . . . . . . . . . . . . . . 20

**9 FRAP experiments to estimate the diffusion coefficient of the LPA-DNA conjugate** **22**

**10 Simulations** **24**

**11 Supplementary movie** **27**

**References** **27**

### 1 Methods

#### 1.1 Reaction assembly and DNA sequences

Reaction was assembled as described in (*1*). Oligonucleotides sequences are displayed in Table S1. Template strands (T and R) were HPLC purified and trigger strands (A) were desalted. For reasons independent of the experiments presented here, minor and inconsequential differences exist between the templates used for the different experiments. These differences are clearly mentioned in the table.

Table S1: DNA sequences with modifications. PTOs are marked with an asterisk * and phosphates with **p**. bt stands for biotin. When biotin-modified oligonucleotides were used, streptavidin was added (200nM in binding sites). Acryd for acrydite modification. For reasons independent of the experiments presented here, minor and inconsequential differences exist between the templates used for the different experiments.

| **Name** | **Sequences 5' → 3'** |
|---|---|
| *Experiment Fig3 MT* | |
| A | TCG TGT TCT GTC |
| T | G*A*C* AGA AC*A CGA GAC AGA ACA C**p** |
| R | A*A*A* GAC AGA ACA CGA**p** |
| *Experiment Fig4 MT* | |
| T | bt-AA GAC AGA AC*A CGA GAC AGA ACA C**p** |
| R | bt-AAA AAA GAC AGA ACA CGA**p** |
| | Acryd-A* A*A*A GAC AGA ACA CGA-Cy5 |
| *Experiment Fig5 MT* | |
| T | G*A*C* AGA ACA CGA GAC AGA ACA C**p** |
| R | A*A*A* GAC AGA ACA CGA**p** |
| *Experiment Fig6 MT* | |
| A | TCG TGT TCT GTC |
| T | G*A*C* AGA AC*A CGA GAC AGA ACA C**p** |
| R | A*A*A* GAC AGA ACA CGA**p** |

#### 1.2 Mechanism of the bistable network

Figure S1 shows a scheme of the bistable network and the details of the three reactions: an autocatalytic reaction of **A** supported by template **T**, a repression of **A** by **R** and a degradation of **A** and **W**. Note **T** and **R** are not degraded by exonuclease as they are 5' protected.

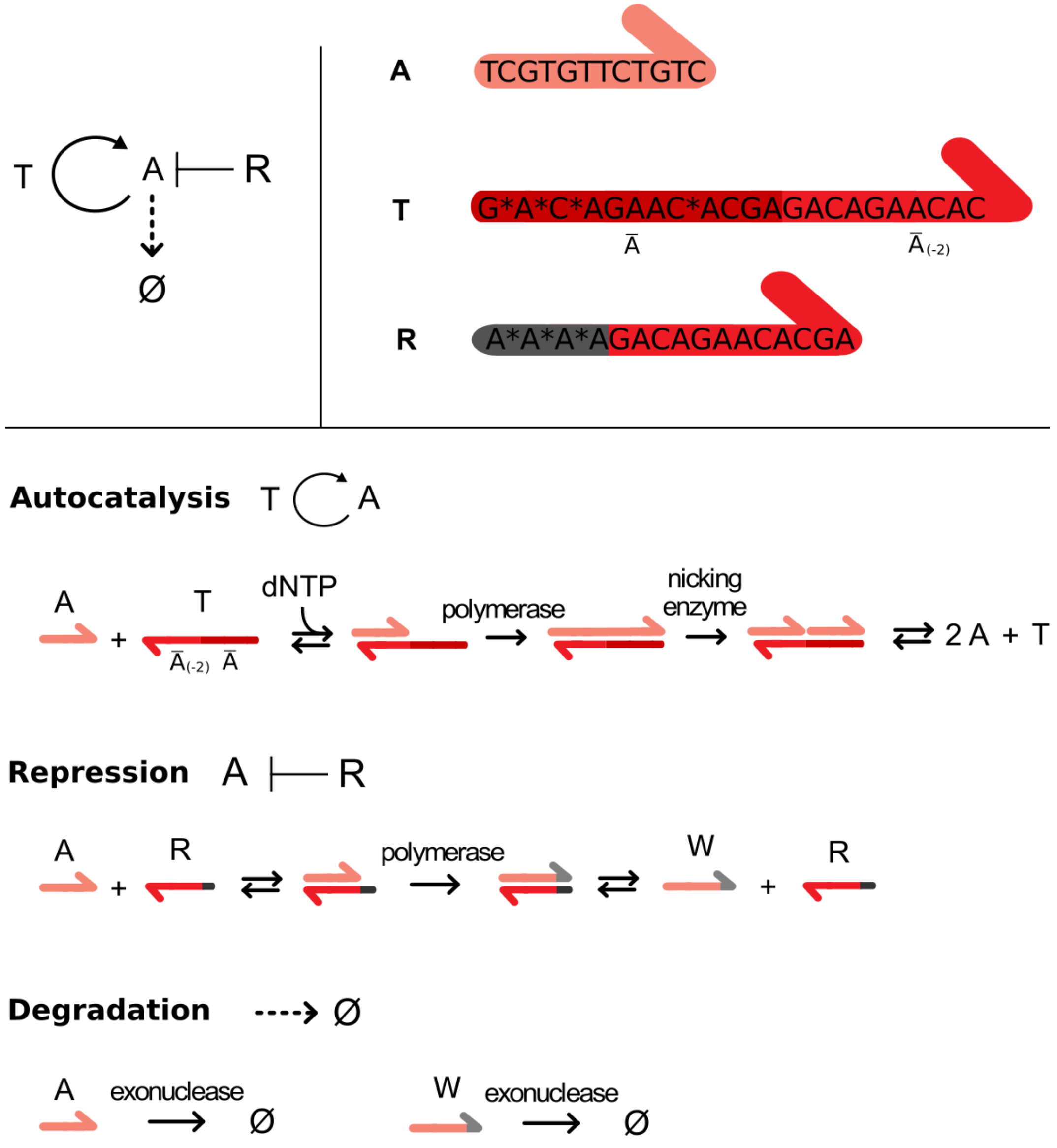

Figure S1: Mechanism of the bistable network

#### 1.3 Synthesis of LPA-DNA conjugate

To synthesize DNA attached to linear polyacrylamide (LPA), we purchase DNA with a 5'-acrydite modification (see scheme of the copolymerization Figure S13a). To start the polymerization reaction of the monomeric acrylamide, tetramethylethylenediamine (TEMED) and ammonium persulfate (APS) are used. Since oxygen can interfere with the reaction, the reaction was done in an argon atmosphere according to the following protocol (Figure S2):

1. Prepare in a Florence flask a 0.5 ml aqueous solution containing 0.12% w/v TEMED, 30 $\mu$M of the acrydite modified DNA and 20% w/w acrylamide. Add a magnetic stirrer.

2. In a second flask, prepare 0.12% w/v APS in water. Only 0.5 ml will be needed, but adding some dead volume is recommended.

3. Close both flasks with septums and use syringe needles to create an argon atmosphere in the flasks. One needle is used to flow argon directly into the solution, another needle is used as outflow for the pressure. This is done for 2 to 3 minutes, then the inflow needles are removed. It is important to use separate sets of needles for the flask, to not accidentally start the reaction.

4. Stir the acrylamide solution described in Step 1 with the magnetic stirrer. Take a syringe, fill it with argon and empty it to remove oxygen. Again, fill in a small volume of argon, suck in 0.5 ml of the APS solution described in Step 2 and suck in some extra argon. Inject the mix into the acrylamide solution under constant stirring. The viscosity should change quickly and the magnet might be stuck.

5. To ensure that there is no oxygen, flow again some argon for 2 to 3 minutes and then use a balloon filled with argon to maintain the argon atmosphere even if there are small leaks. Let the reaction proceed over night.

6. On the next day, the reaction is done. To get the magnetic stirrer to work better, 1 ml water is added. If the magnet is stuck, it can help to use a second magnet to move it a bit. Add approximately 2 ml acetone. A visible blob should form.

7. Use a Büchner funnel to remove most of the acetone and compounds which did not react. Then vacuum dry the pellet for several hours. When it is dry, it should weigh 100 mg.

8. Dissolve the pellet in 5 ml water (a rotator was used, it takes several hours). This will give a final concentration of 2% w/w LPA. Higher concentrations are hard to pipette due to the high viscosity. The expected DNA concentration is 3 $\mu$M.

The final solution might contain unreacted acrydite strands or strands with only small LPA attachements, for example due to cyclization. We use electro elution to remove those strands. For electroelution we use a lab-built device, with a 50 mm wide chamber that is sandwiched between two porous membranes with 0.22 $\mu$m diameter holes. 98 $\mu$l of the LPA-DNA solution is mixed with 2 $\mu$l 50xTAE buffer and filled into the chamber. Each membranes separates the chamber from a tank containing 3.5 ml 1xTAE buffer. We put an electrode into each tank and run a current of 3 mA, resulting in a voltage of approximately 50-80 V. After electro elution, the TAE buffer is exchanged with Amicon centrifugal filters Ultracel (10 K). Since it is expected that DNA is removed during electro elution, the volume is not brought back to the initial volume. Water is added until a consistency is reached that allows pipetting without problems. In the end, the concentration of DNA is measured with a nanodrop device. Since LPA also absorbs at a wavelength of 260 nm a pure LPA reference sample is used.

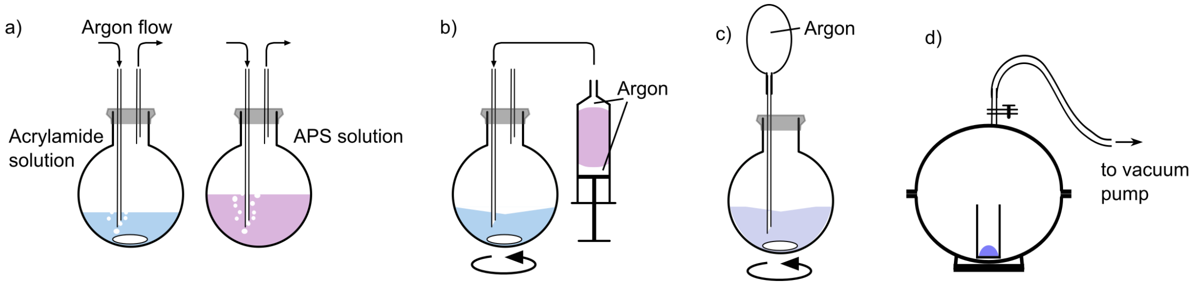

Figure S 2: Synthesis of LPA-DNA conjugate. a) TEMED, monomeric acrylamide and acrydite modified DNA solution and APS solution are prepared separately and maintained in an argon atmosphere. b) Solutions are mixed under argon atmosphere. c) Overnight reaction. d) Vacuum drying to get a LPA-DNA conjugate pellet.

#### 1.4 Microscopy imaging

DNA concentration over space and time was measured by fluorescence. The initial gradient was visualized through a Cy5 dye attached to the 3' end of the strand. The DNA production induced by the bistable program was recorded by the green fluorescence from EvaGreen dye (1x EvaGreen DNA binder, 20x dilution of the manufacturer's stock solution, Biotium). In this case, the fluorescence signal is proportional to the concentration of double stranded DNA. Once the gradient was formed, the capillary (or the gel) was dropped into a pool of mineral oil (Sigma) to keep the capillary at constant temperature and avoid evaporation. The pool was made out of a 5 mm thick plastic frame sealed on top of a glass slide using Araldite epoxy glue. The fluorescence along the capillary was recorded on a Zeiss Axio Observer Z1 fully automated epifluorescence microscope equipped with a CoolLED pE-2 fiber-coupled illuminator, the corresponding filter sets, a Marzhauser XY motorized stage, a Tokai Hit thermo plate, and an Andor iXon Ultra 897 EMCCD camera. These instruments were controlled with MicroManager 1.4. Contiguous images were recorded automatically every 1 to 20 minutes using a 2.5 $\times$ objective to get an entire view of the capillaries after stitching. The kymographs were obtained by averaging the fluorescence images over the width of the capillary and then stacking these profiles over time.

#### 1.5 Gradient generation protocol

To create a gradient of DNA strand **R** in a homogeneous solution of all other species, two solutions were prepared, one with **R** and the other without. A micropipette was connected to a rectangular glass capillary (Vitrocom) through a PDMS connector, as depicted in Figure S3. The capillary was first filled with the solution containing no **R**. Subsequently, the end of the capillary was dipped into the reaction solution containing **R**. Micropipette up-and-down pumps of a volume in the order of a quarter of the total volume of the capillary were performed to create the gradient by Taylor dispersion.

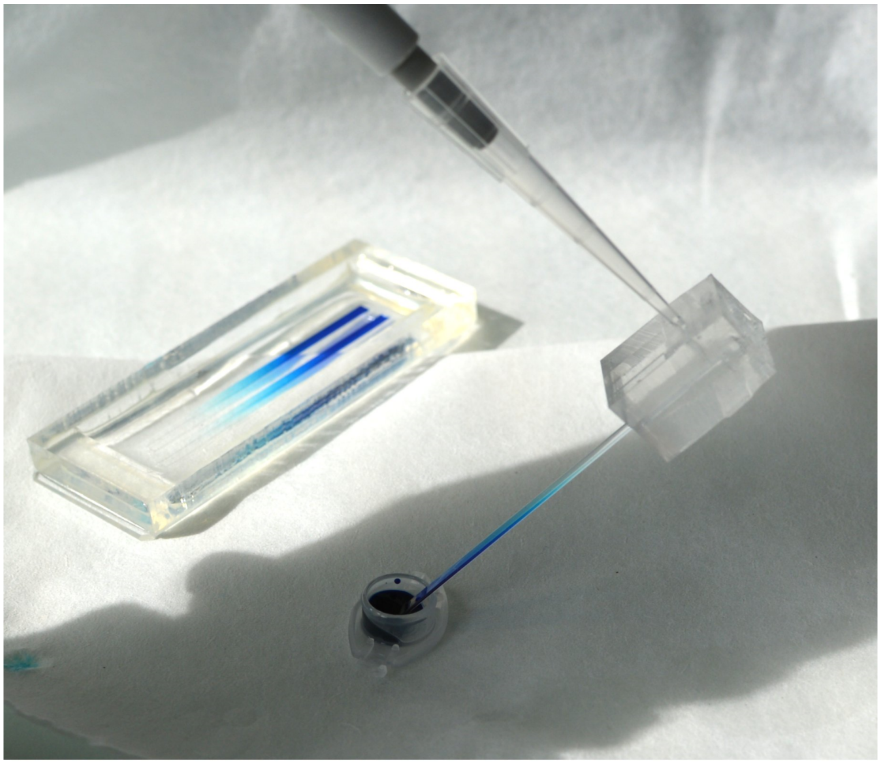

Figure S3: Photograph of the setup used to generate a gradient inside a capillary through back and forth pipetting.

The reusable PDMS connector was prepared as follows. A sacrificial capillary is treated with Trichloro(1H,1H,2H,2H-perfluorooctyl)silane antiadhesive (Sigma) before being vertically poured in a 1 cm thick freshly prepared PDMS. The capillary must be kept 2 mm above the bottom of the tank. After PDMS cross-link, the capillary is gently removed, the PDMS is cut by hand if necessary and a 1 mm in diameter hole is finally punched to later connect the pipette tip.

Typically, for a $0.2 \times 2 \times 50$ mm 20 $\mu$L capillary, 15 up-and-down pumps of 6 $\mu$L were performed, for a $0.2 \times 4 \times 50$ mm 40 $\mu$L capillary, 15 up-and-down pumps of 12 $\mu$L were performed, and for a $1 \times 1 \times 50$ mm 50 $\mu$L capillary, 15 up-and-down pumps of 15 $\mu$L were performed. More pumping cycles are needed when DNA is bound to LPA.

As illustrated in Figure S4, 30 up-and-down pumps allows to get a smooth gradient of DNA but, because of a reduced diffusion, the gradient is less smooth when DNA is bound to LPA. The slow diffusion of DNA-LPA conjugate makes it difficult to prepare a smooth gradient by hand as more than 50 up-and-down pumps are needed. For convenience, instead of using a manual pipette, we automatized the gradient generation using a programmable syringe pump (Nemesys, CETONI GmbH, Germany) and the same PDMS connector. Supplementary Movie S1 shows the automated preparation of a methylene blue dye gradient in a $2 \times 0.2 \times 50$ mm capillary.

The same protocol was used to mold low gelling agarose (Sigma A9539, low EEO) with embedded DNA/enzyme solutions and underlying DNA gradient as detailed in the main text. 2% agarose gel diluted in 2x concentrated active solution were typically used and mixed in a 1/1 ratio before being used to prepare the gradient. $1 \times 1 \times 50$ mm capillaries were used to finally get pieces of gel that can be easily handled by hand.

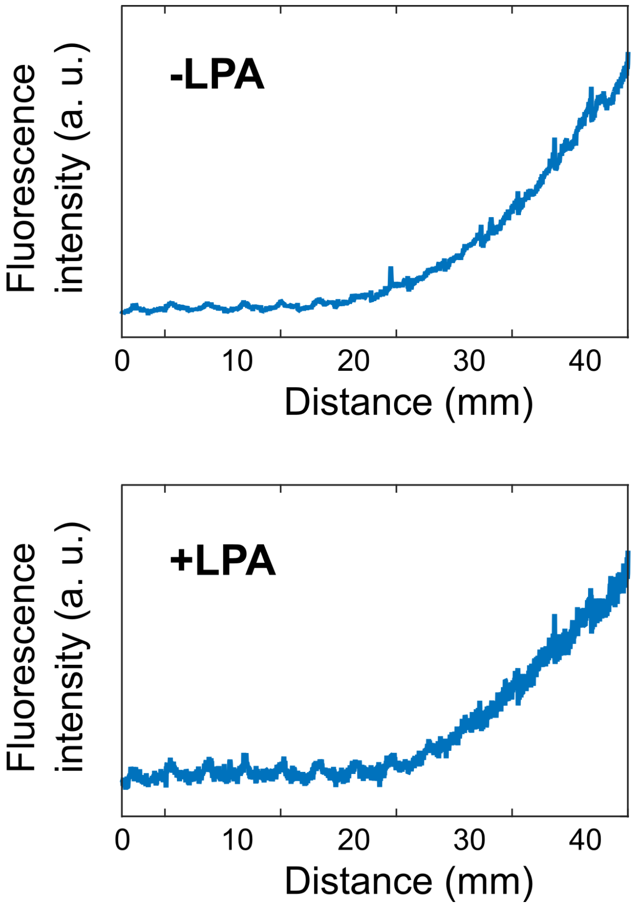

Figure S4: Fluorescence intensity profiles of gradients obtained with 30 up-and-down pumps with DNA and DNA attached to LPA.

### 2 Stability of the enzymes after temperature perturbation

Experiments performed in agarose gel require heating the DNA/enzyme solution to 60°C during a few minutes to keep the gel liquid during the loading and the gradient preparation.

We designed three experiments to test the enzyme activity following an incubation at high temperature: replication, degradation and nicking assays.

The replication experiment consists of adding 100 nM of DNA template T to a preincubated solution that contains polymerase and nicking enzyme. The degradation assay consists of adding 1 $\mu$M of an unprotected 16-mer DNA to a preincubated solution that contains exonuclease. The nicking enzyme assay consists of adding 1uM of a specific hairpin DNA with a dye-quencher modification -

FAM-CGCTCGTGGATCCAGAAAAAAAAAACTGGATCCACGAGCG-Dabcyl, that will be nicked to release a fluorescent strand - to a preincubated solution that contains nicking enzyme.

As shown Figure S5, after a 17 hours incubation at 61°C, replication, degradation and nicking dynamics are similar to the experiments without incubation. We confirmed the enzyme activity is only affected for very long incubation time, and that gel preparation requirements - a few minutes - doesn't affect the DNA toolbox.

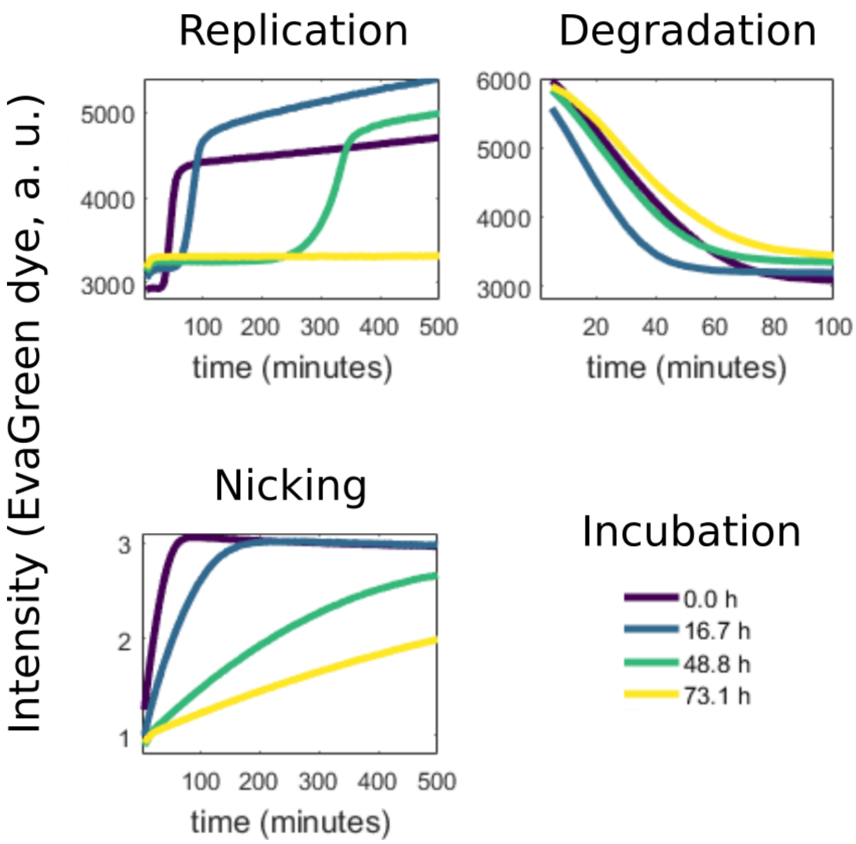

Figure S5: Stability of the enzyme activity after temperature perturbation for gel sample preparation. Experiments are performed at 44°C after incubation at 61°C for 0 hour, 17 hours, 49 hours and 73 hours.

### 3 Parasite suppression strategy

Our method to prevent the formation of parasite strands in an exponential amplification reaction by suppressing untemplated replication relies on specific nicking enzymes such as Nb.BssSI that bears only three types of bases on each of the strands of the recognition site.

As shown in Figure S6, fully functional replication templates (template **T**) can be designed using A C G bases only, that will produce DNA strands containing T G C bases only (product **A**). In the absence of dATP in the buffer, any parasitic strands formed will not contain the A base. They will therefore not carry the nicking enzyme recognition site and will not be replicated exponentially, unlike the replicate **A**.

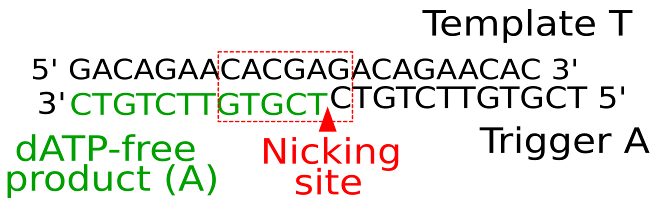

Figure S6: Parasite suppression strategy. Details of the Nb.BssSI nicking enzyme recognition site.

### 4 Quantification of evaporation

**In solution** - We estimated evaporation by measuring the volume of mineral oil entering the capillary during a 7-day experiment at 44°C. After 7 days, 1.5% of the capillary was evaporated.

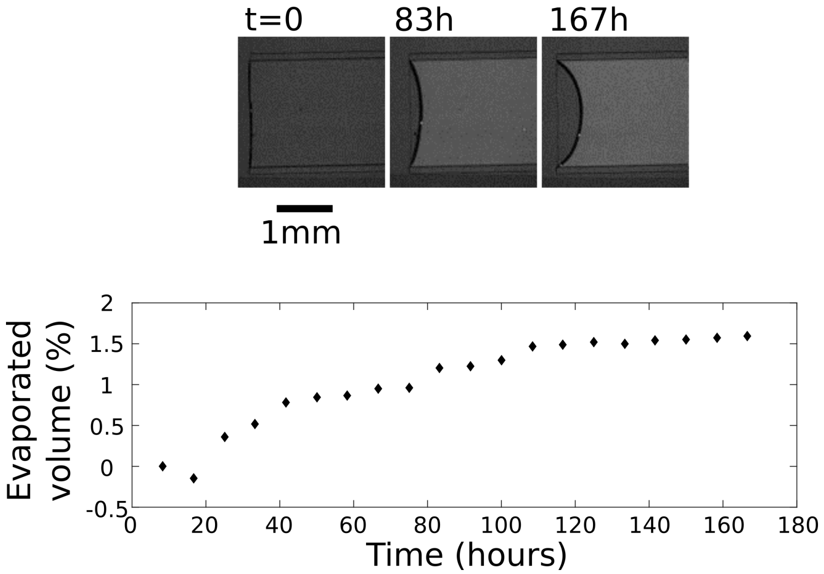

Figure S7: Evaporation during a 7-day experiment at 44°C. The fluorescence intensity inside the capillary increases not because of evaporation but because of the diffusion of a fluorescent DNA gradient.

**In gel** - We estimated evaporation by measuring the surface of a gel during a 32 hours experiment at 44°C. From the surface measurements and considering an isotropic contraction, we found the volume of the gel was linearly decreasing by 14% in 24 hours.

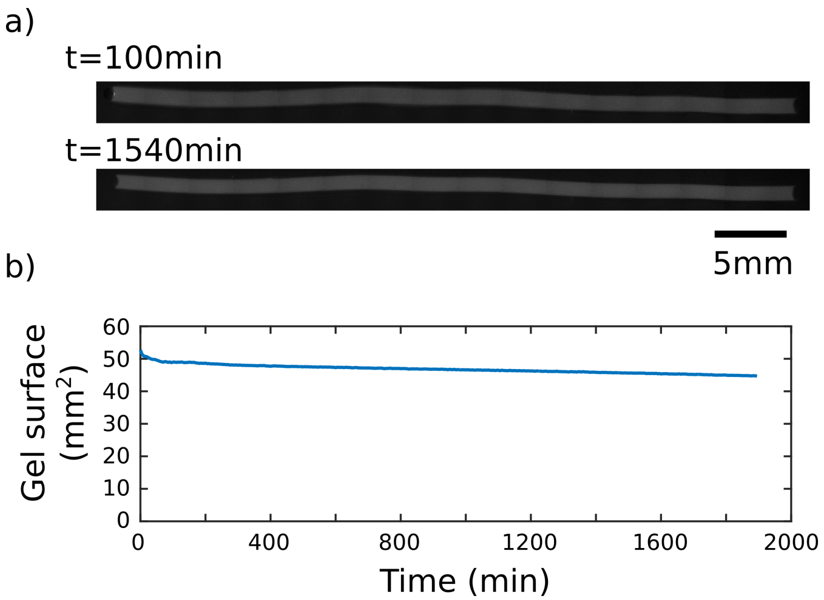

Figure S8: Evaporation in an agarose gel: the gel contracts over time, which can be analyzed by fluorescence microscopy and image thresholding. a) Pictures of gel showing contraction after 24 hours at 44°C in a oil pool. b) Gel surface as a function of time.

### 5 Parasite in 1D

Figure S9, corresponding to Figure 3a left kymograph in the main text, shows the parasite appearance. We have noticed that parasites form preferentially near defects in the channel (dust, extremities, air bubbles) and generally appear in several positions, almost simultaneously.

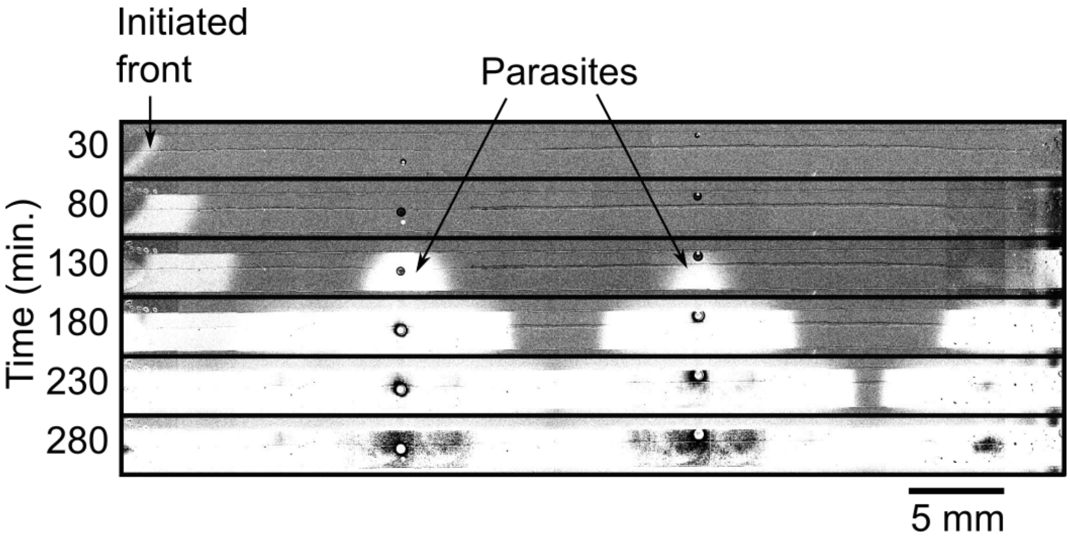

Figure S9: A reaction-diffusion front disrupted by the exponential amplification of a DNA parasite strand.

### 6 Front concentration profiles

Figure S10, corresponding to Figure 3 in the main text, shows the profiles of fluorescence intensity along the channel, with 100 min intervals, for both traveling front and two-band pattern experiments.

The fluorescence of EvaGreen is proportional to the concentration of dsDNA in the capillary. In the presence of a gradient of repressor (Figure S10b), it induces an increase in fluorescence at the front head during propagation as produced **A** strands bind to an increasing concentration of **R**.

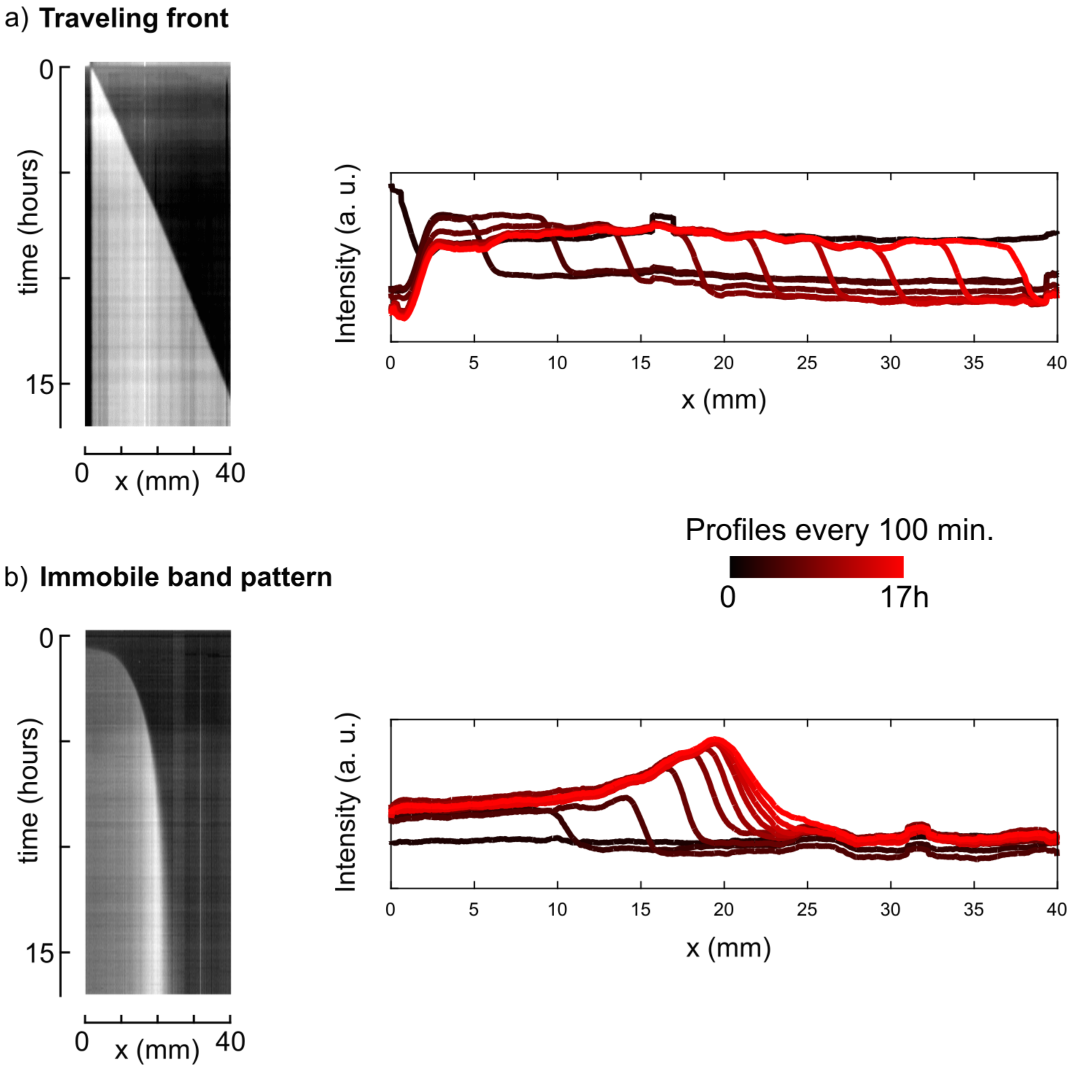

Figure S10: Kymographs and profiles of fluorescence intensity along the capillaries, in 100 min. intervals, for (a) a traveling front experiment and (b) a two-band pattern experiment.

### 7 Underlying gradient diffusion with and without LPA

The experiment in Figure 4a in the main text shows that the stability of the band pattern is not maintained if the underlying gradient DNA strands are not copolymerized with linear polyacrylamide chains.

Figure S11 shows the diffusion of the gradient with and without LPA attached to the DNA after 10 minutes, 50 hours and 125 hours. Note the gradient recorded here is similar, but doesn't correspond to the experiment in the main text as fluorescence images of both the gradient and the produced DNA are difficult to acquire simultaneously.

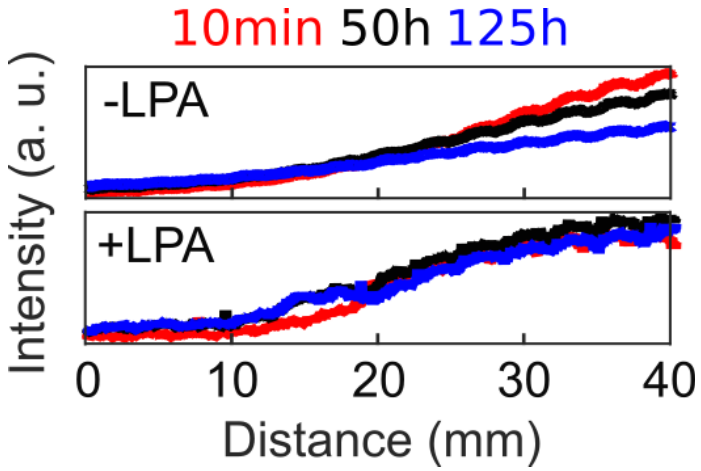

Figure S11: Concentration profiles of repressor gradient copolymerized or not with linear polyacrylamide chains after 10 minutes, 50 hours and 125 hours.

### 8 Estimation of the molecular weight distribution of the LPA-DNA conjugate

#### 8.1 By size exclusion chormatography

We used size exclusion chromatography (SEC) to estimate the weight distribution of the LPA-DNA conjugate. SEC was performed using a Superdex 200 Increase 10/300 GL column (GE) in water. We injected 150 $\mu$L of 1.5 $\mu$M electroeluted LPA-DNA at 0.5 mL/min. For comparison, we performed subsequent injections of commercial samples of LPA, one being sold as $M_n = 0.1 - 0.2$ MDa (Aldrich #179222) and the second one as $M_w = 5 - 6$ MDa (Aldrich #92560). The eluted sample was detected by absorbance at 190 nm. Figure S12 shows that all three samples had one main peak in a similar range of elution volumes: $7.1 - 10.8$, $7 - 11.3$ and $6.8 - 9$ mL, for LPA-DNA, $M_n = 0.1 - 0.2$ LPA and $M_w = 5 - 6$ MDa LPA, respectively. However, lower weight LPA displayed a smaller peak at higher elution volumes (smaller $M_w$) while the higher weight LPA had a broad peak at very low elution volumes (high molecular weights) and the LPA-DNA displayed a single rectangular peak. We concluded that the weight distribution of the commercial samples is too wide and difficult to rely to the $M_n$ and $M_w$ values provided by the manufacturer to serve as molecular weight standards. To estimate the $M_w$ of LPA-DNA we thus relied on the theoretical calibration curve of elution volume $V_{el}$ *vs.* molecular weight $M_w$, $log(M_w) = -0.2V_{el} + 7.8$, provided by the manufacturer of the column. With this we conclude that the LPA-DNA conjugate that we synthesized had a molecular weight in the range $0.3 - 2$ MDa.

#### 8.2 From the diffusion coefficient

Scholtan measured the diffusion coefficients of linear polyacrylamide as a function of molecular weight in the range $20 - 500$ kDa, and obtained the relation $D = 8.46 \times 10^4 M_w^{-0.69}$ $\mu$m$^2$/s, with $M_w$ being expressed in g/mol, in water at 20 °C. In the MT we measured $D =$

$4.8 \pm 2\ \mu m^2/s$ for the LPA-R copolymer at 40 °C. After correcting from the temperature variation of $D$, this value corresponds to $M_w = 1.4 - 2.5$ MDa, which is consistent with the SEC estimation.

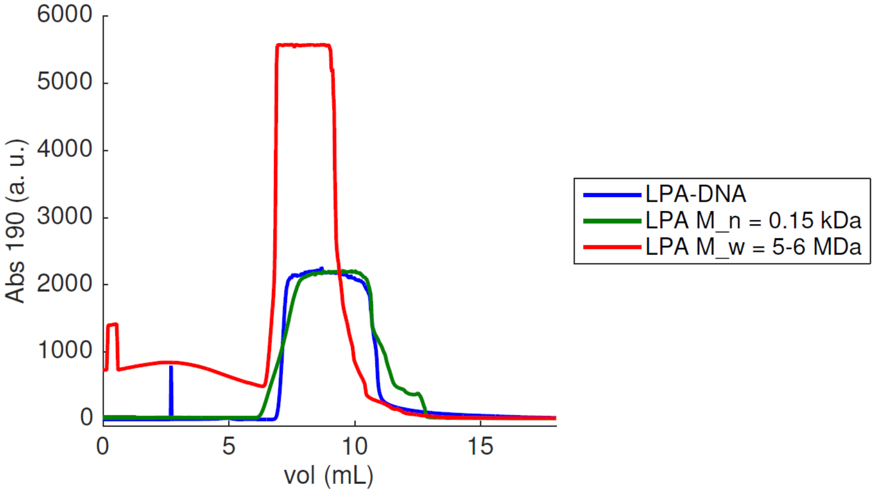

Figure S12: Size exclusion chromatograms of LPA-DNA conjugate compared with commercial LPA samples.

### 9 FRAP experiments to estimate the diffusion coefficient of the LPA-DNA conjugate

We estimated the diffusion coefficient of DNA and LPA-DNA conjugate using Fluorescence Recovery after Photobleaching (FRAP) experiments. FAM labelled-DNA with acrydite modification was used,

Sequences 5' → 3': Acryd-CATTCGGCCGAATG-FAM

FRAP experiments were performed using a standard microscope, a 20x objective was used for bleaching and a 2.5x objective for recording the fluorescence recovery. 0.2*4*50mm capillaries were filled with DNA or LPA-DNA and setup onto the microscope heating stage. Samples were heated to 40°C. A 30s bleaching was first done, then a 20 to 45s timelapse was performed to record the fluorescence recovery.

The image stacks were further analyzed by FFT using Matlab to determine the diffusion coefficient of the DNA and the LPA-DNA (Figure S13 panel b).

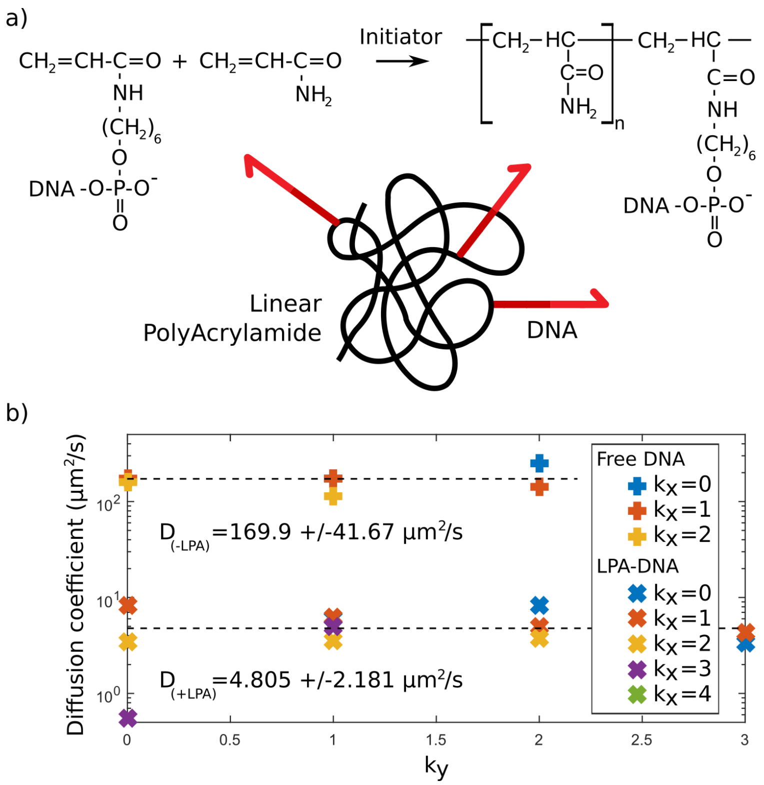

Figure S13: Reducing the diffusion of DNA by attaching a long linear polyacrylamide chain to the strand. a) Scheme of the copolymerization of 5'-acrydite-modified DNA into polyacrylamide. Monomeric acrylamide, tetramethylethylenediamine (TEMED) and ammonium persulfate (APS) were used to polymerize linear acrylamide in the presence of modified DNA. Unreacted acrydite DNA strands or strands attached to short polymer chains were removed by electro-elution. b) FRAP (Fluorescence Recovery After Photobleaching) was used to estimate the diffusion coefficient of a free DNA and a LPA-DNA complex. Diffusion coefficients were determined by FFT analysis using MATLAB. Experiments were performed at 40 °C.

### 10 Simulations

Matlab simulations of a simplified model capture the consequence of a diffusing, underlying gradient on the long-term stability of the two-band pattern. It takes into account hybridization kinetics, production and degradation pathways of strand **A** and waste **W** as well as the diffusion of **A**, **W**, template **T** and repressor **R**. In the model, possible hybridization of **W** to **T** is also taken into account.

In the model, the following enzymatic steps are considered. **A** produces more **A** with reaction rate constant **r** only when bound to **T**. Since the concentration of **T** is constant, the reaction rate of **A**-production is limited. The conversion of **A** (when bound to **R**) to **W** takes place with the same reaction rate constant **r**. For the enzymatic degradation of **A** and **W**, we assume exponential degradation as was done in a simple DNA-toolbox model elsewhere (*2*).

We used the Matlab built-in PDE solver 'pdepe'. Our model consists of eight equations, where $X_Y$ means X bounds to Y

$$\frac{\partial A}{\partial t} = -k_{on} \cdot A \cdot R + k_{off}^{A} \cdot R_A - k_{on} \cdot A \cdot T + k_{off}^{A} \cdot T_A + r \cdot T_A - d \cdot A + D_A \cdot \frac{\partial^2 A}{\partial x^2}$$

$$\frac{\partial T}{\partial t} = -k_{on} \cdot A \cdot T + k_{off}^{A} \cdot T_A - k_{on} \cdot W \cdot T + k_{off}^{A} \cdot T_W + D_T \cdot \frac{\partial^2 T}{\partial x^2}$$

$$\frac{\partial T_A}{\partial t} = +k_{on} \cdot A \cdot T - k_{off}^{A} \cdot T_A + D_T \cdot \frac{\partial^2 T_A}{\partial x^2}$$

$$\frac{\partial W}{\partial t} = -d \cdot W - k_{on} \cdot W \cdot T + k_{off}^{A} \cdot T_W - k_{on} \cdot W \cdot R + k_{off}^{W} \cdot R_W + D_A \cdot \frac{\partial^2 W}{\partial x^2}$$

$$\frac{\partial T_W}{\partial t} = +k_{on} \cdot W \cdot T - k_{off}^{A} \cdot T_W + D_T \cdot \frac{\partial^2 T_W}{\partial x^2}$$

$$\frac{\partial R}{\partial t} = -k_{on} \cdot W \cdot R + k_{off}^{W} \cdot R_W - k_{on} \cdot A \cdot R + k_{off}^{A} \cdot R_A + D_R \cdot \frac{\partial^2 R}{\partial x^2}$$

$$\frac{\partial R_A}{\partial t} = +k_{on} \cdot A \cdot R - k_{off}^{A} \cdot R_A - r \cdot R_A + D_R \cdot \frac{\partial^2 R_A}{\partial x^2}$$

$$\frac{\partial R_W}{\partial t} = +k_{on} \cdot W \cdot R - k_{off}^{W} \cdot R_W + r \cdot R_A + D_R \cdot \frac{\partial^2 R_W}{\partial x^2}$$

Most parameters were extracted from previous experiments and publications, $d$ and $k_{off}^{W}$ were adjusted by hand. The initial gradient is a polynomial fit of an experimental fluorescent DNA gradient. Since selfstart is not part of the model, a homogenous distribution of 1 nM **A** was used as initial condition.

$D_A = 1.23 \cdot 10^{-2}\, mm^2 \cdot min^{-1}$

$D_T = 7.20 \cdot 10^{-3}\, mm^2 \cdot min^{-1}$

$D_R = 3.00 \cdot 10^{-4}\, mm^2 \cdot min^{-1}$ when bound to LPA or $1.00 \cdot 10^{-2}\, mm^2 \cdot min^{-1}$ when unbound.

$kon = 0.06\, nM^{-1} \cdot min^{-1}$

$k_{off}^{A} = 0.18\, min^{-1}$

$k_{off}^{W} = k_{off}^{A}/30\, min^{-1}$

$r = 0.2\, min^{-1}$

$d = 5.0\, min^{-1}$

The model reduces the complexity of the experimental system in several ways:

1. In the model, **T** only binds a single **A**.

2. Dehybridization of **T** and **A** is described by a single off-rate $k_{off}^{A}$. This does not take into account that the template is asymmetric and that the $k_{off}$ rate on the 5'-end of the template changes when strand displacement comes into play.

3. **A** bound to **T** produces free **A** in a one-step process. Experimentally, this step consists of elongation, nicking and dehybridization/strand displacement.

4. For the creation of **A** from $\mathbf{T_A}$ and the conversion of $\mathbf{R_A}$ to $\mathbf{R_W}$, the same rate **r** is used. The second reaction should in principle be faster, since it is a one-step process facilitated by the polymerase, whereas the first reaction consists of the three steps mentioned above.

5. Degradation by exonuclease does not saturate.

The described differences do not constitute an exhaustive list, but include reactions that could be implemented rather easily at the cost of adding complexity to the model. Other effects that are harder to capture are surface effects, change of enzymatic function over time due to changed dNTP and pyrophosphate concentration or denaturation of enzymes in the long-term experiments. In addition, when **A** is bound to the 3'-end of **T**, it has two unbound nucleotides that might be cleaved off by exonuclease, resulting in changes in $k^{A}_{off}$ and $k^{W}_{off}$.

The full code is available upon request.

#### 11 Supplementary movie

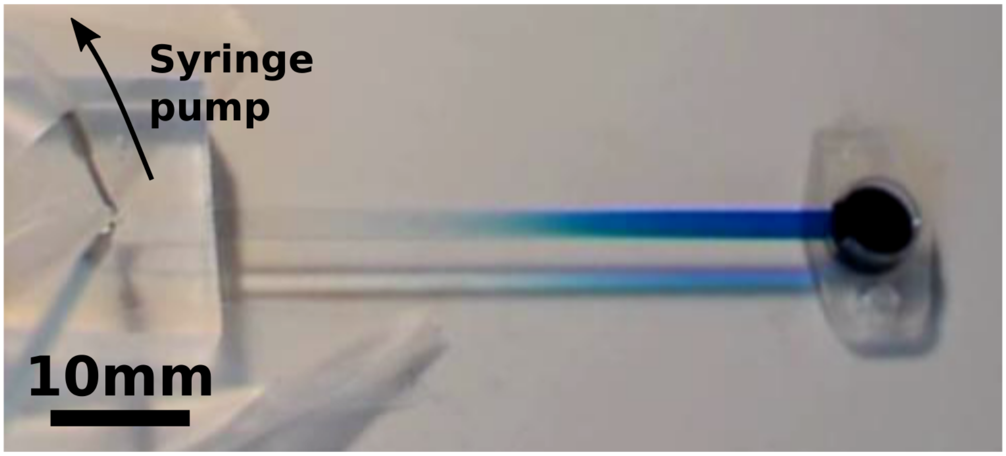

Movie S1: Automated generation of the gradient using a programmable syringe pump. Methylene blue dye was used for visualization purpose. The movie is real time.
